# Contrast-enhanced ultrasound imaging detects anatomical and functional changes in rat cervical spine microvasculature with normal aging

**DOI:** 10.1101/2024.03.12.584672

**Authors:** Jennifer N. Harmon, Preeja Chandran, Abarajithan Chandrasekaran, Jeffrey E. Hyde, Gustavo J. Hernandez, May J. Reed, Matthew F. Bruce, Zin Z. Khaing

**Affiliations:** Department of Neurological Surgery, University of Washington, 1959 NE Pacific St., Seattle, WA, USA; Department of Gerontology and Geriatric Medicine, University of Washington, Seattle, WA, USA; Applied Physics Laboratory, University of Washington, Seattle, WA, USA

**Keywords:** Cervical spinal cord, perfusion, hypoxia, pericytes

## Abstract

Normal aging is associated with significant deleterious cerebrovascular changes; these have been implicated in disease pathogenesis and increased susceptibility to ischemic injury. While these changes are well documented in the brain, few studies have been conducted in the spinal cord. Here, we utilize specialized contrast-enhanced ultrasound (CEUS) imaging to investigate age-related changes in cervical spinal vascular anatomy and hemodynamics in male Fisher 344 rats, a common strain in aging research. Aged rats (24-26 mo., N=6) exhibited significant tortuosity in the anterior spinal artery and elevated vascular resistance compared to adults (4-6 mo., N=6; tortuosity index 2.20±0.15 vs 4.74±0.45, p<0.05). Baseline blood volume was lower in both larger vessels and the microcirculation in the aged cohort, specifically in white matter (4.44e14±1.37e13 vs 3.66e14±2.64e13 CEUS bolus AUC, p<0.05). To elucidate functional differences, animals were exposed to a hypoxia challenge; whereas adult rats exhibited significant functional hyperemia in both gray and white matter (GM: 1.13±0.10-fold change from normoxia, p<0.05; WM: 1.16±0.13, p<0.05), aged rats showed no response. Immunohistochemistry revealed reduced pericyte coverage and activated microglia behavior in aged rats, which may partially explain the lack of vascular response. This study provides the first *in vivo* description of age-related hemodynamic differences in the cervical spinal cord.

## Introduction

Vascular changes associated with aging are directly implicated in the pathogenesis of neurodegenerative and cognitive disorders (e.g., Alzheimer’s disease, vascular dementia) and are thought to predispose aged individuals to more severe cerebral ischemic injuries (e.g., stroke) (1–5). Previous clinical findings have shown worse functional outcomes in aged patients following spinal cord injury (SCI), suggesting that the aged spinal cord similarly exhibits enhanced susceptibility to ischemic injury (6). However, most existing research focuses on brain vasculature; there are relatively few studies that describe age-related changes in spinal vasculature, despite the progressive increase in the mean age of spinal cord injured patients (7, 8). To our knowledge, only one major study has explicitly investigated vascular differences in the spinal cord due to normal aging; the authors found that aged animals exhibited enhanced tortuosity and reduced vessel density compared to young and adult animals (9). Additional recent studies have investigated age-specific vascular contributions to progressive degeneration following SCI and vascular disruption in response to hypoxia exposure, but these studies relied solely on histological or *ex vivo* imaging methods and did not elucidate underlying baseline vascular differences that may render the aged cord more susceptible to injury (10, 11). There is a dearth of evidence elucidating vascular changes in the spinal cord with normal aging *in vivo*, particularly with respect to hemodynamic and functional deficits.

Most *in vivo* studies of spinal hemodynamics have relied on optical imaging techniques, such as laser speckle contrast imaging or two-photon microscopy (12, 13). While these methods provide excellent resolution, they are severely limited in imaging depth due to light scattering and absorption (14, 15). Ultrasound, on the other hand, is capable of imaging the entire depth of the spinal cord with excellent spatiotemporal resolution (∼ 100 um, up to 20 kHz). Recent advances in specialized contrast-enhanced ultrasound (CEUS) modalities have enabled both super-resolution imaging of spinal vascular anatomy and sensitive quantification of microcirculatory hemodynamics (16, 17). While ultrasound localization microscopy (ULM) has been applied to study age-related changes in brain microvasculature, the spinal cord remains unstudied (18).

Here, we utilized ultrasound imaging to study anatomical and functional differences in cervical spinal vasculature with normal aging. CEUS and ULM revealed significant alterations in both tissue and vascular anatomy with normal aging. In conjunction with these findings, multiple modalities identified increased vascular resistance in the aged spinal cord, as well as a significant reduction in blood volume at baseline. To elucidate functional deficits in the aged spinal microvasculature, animals were subjected to a hypoxia challenge; while adult animals exhibited hyperemia in both gray and white matter, aged animals showed no response whatsoever. Histological analysis was conducted to establish a potential mechanism for this discrepancy; both pericyte and microglia behavior throughout the spinal cord were dysregulated. Specifically, we observed a reduction in pericyte coverage of microvessels and an increase in activated microglia. The current study provides the first comprehensive evaluation of age-dependent changes in spinal hemodynamics and provides a basis for understanding increased susceptibility to ischemic injury in the aged spinal cord.

## Materials and methods

### Animal model

All animal procedures were approved by the Institutional Animal Care and Use Committee (IACUC) at the University of Washington (Protocol #4362-01). Experiments were conducted in accordance with NIH Office of Laboratory Animal Welfare (OLAW) ARRIVE guidelines.

Adult (4-6 mo., N=6) and aged (24-26 mo., N=6) male Fisher 344 rats were acquired from the National Institute on Aging (Bethesda, MD). Rats were anesthetized with isoflurane (5% induction, 1-3% maintenance) prior to tail vein catheterization (24 Ga). The catheter was flushed with heparinized saline (1%, 0.5 mL) and fitted with a three-way valve for administering bolus injections of microbubble contrast agent (Definity, Lantheus, Billerica, MA) during scans. Bubble dosing was adjusted (215 µl/kg followed by 0.2 mL saline flush) to account for significant differences in body weight between adult and aged animals (351±4.91g vs 517±14.9g, p<0.05). Topical analgesic (1.5 mg/kg Lidocaine, 1 mg/kg Bupivacaine) was administered following shaving and sterilization of the area above the C2-T2 segments. A #10 scalpel blade was used to make a 2.5 cm longitudinal incision centered over C4-C6. Following subperiosteal dissection of paraspinal muscles, a three-level laminectomy was performed to expose levels C4-C6 for epidural ultrasound imaging. Animals were stabilized at the C2 and T2 spinous processes using a custom frame to minimize motion during imaging sessions.

### Experimental design and ultrasound imaging

All imaging sequences were implemented in-house on a research ultrasound system (Vantage 128, Verasonics, Kirkland, WA) using a 15 MHz linear array transducer (L22-14vX, Verasonics). Warmed sterile saline and ultrasound gel (Aquasonic) were applied to the exposed spinal cord for acoustic coupling. An initial set of baseline scans was conducted longitudinally with the transducer aligned at midline. The transducer was then rotated 90°, aligned to the center of C5, and a set of axial baseline scans was acquired. Each set of scans consisted of: (1) a 3D stepped B-mode sequence (6 mm rostrocaudal extent, 25 µm step size); (2) plane wave Doppler imaging; (3) plane wave contrast-enhanced ultrasound (CEUS) imaging during a bolus injection of contrast; (4) sparse CEUS acquisitions for ultrasound localization microscopy (ULM); (5) a 3D stepped CEUS sequence (6 mm rostrocaudal extent, 25 µm step size). The development and implementation of the imaging setup and sequences have been described in detail in previous publications from our group (16, 17, 19, 20). Following baseline scans, animals were subjected to hypoxia as a vascular challenge. CEUS bolus imaging (32 sec total acquisition time) was initiated 90 sec after switching from 20% O_2_ to 10% O_2_ (balanced in N_2_; Airgas, Radnor, PA). Gases were switched back after 3 min of total hypoxia exposure. Hypoxia bolus scans were all oriented axially and aligned at C5. Following terminal imaging, animals were injected with biotinylated lectin (2 µl, B-1175-1, Vector Laboratories, Newark, CA) and subsequently euthanized using an overdose of sodium pentobarbital (Euthasol, IP, ≥50 mg/kg).

### Tissue volume measurements

The gray matter (GM), white matter (WM), and overall spinal cord parenchyma volumes were measured using a custom Matlab script (R2019b, Mathworks, Natick, MA). The total cord parenchyma volume was measured from 3D B-mode acquisitions. The GM and WM volumes were measured using 3D CEUS acquisitions. In brief, 3D CEUS scans consisted of short (40 frame) contrast-specific plane wave ensembles (667 Hz compounded pulse repetition frequency; PRF) acquired per spatial step (25 µm). These were used to generate a power Doppler image for each location; the Doppler power varies proportionately to local blood volume, resulting in visible contrast between gray and white matter. Masks were drawn over the full cord and GM to calculate volume for GM and WM individually.

### Ultrasound localization microscopy

Long dwell-time CEUS acquisitions (720 frames, 1.8 sec, 400 Hz compounded PRF) were used for ULM (16, 17). Acquisition was initiated 4-6 mins after a bolus injection of contrast agent to achieve a sparse concentration of microbubbles within the vasculature. Data were wall filtered using the singular value decomposition (SVD) to isolate signal from larger vessels (projections 125-500) prior to microbubble localization and tracking across successive frames. Localization was achieved using peak-identification following cross-correlation with an estimated point spread function, and tracking was done using a partial assignment method based on the Kuhn-Munkres algorithm (16, 21). Tracks were smoothed using a Savitsky-Golay filter (first order, 3 frame window) and linearly interpolated by a factor of 10. Tracks shorter than 7 frames were excluded to reduce noise. Track data were accumulated across a total of 15-20 acquisitions (27-36 sec) to generate a single composite ULM image. Longitudinal ULM images were used to calculate vessel tortuosity using established methods (22).

### Bolus kinetics

Contrast-specific plane wave ensembles (40 frames, 667 Hz PRF) were acquired at a 16 Hz inter-ensemble acquisition rate. Imaging was initiated immediately prior to contrast injection and lasted for a total of 32 seconds to capture a full dynamic bolus inflow. Each ensemble was filtered using the SVD to remove corrupting signal from larger vessels (retained projections 1-5) prior to generating a power Doppler image. A lognormal curve fitting routine was applied to time-intensity data on a pixelwise basis to generate spatial maps of bolus inflow parameters. The area under the curve (AUC) and the rise time (RT; time from 5% of peak intensity to 95% of peak intensity) were calculated. AUC is proportional to local blood volume, whereas RT is analogous to vascular resistance. Parametric maps were analyzed using a custom Matlab script. For AUC analysis, separate masks were drawn over GM and WM as described for tissue volume measurements.

### Doppler imaging

A 5-angle plane wave sequence (4 kHz compounded PRF) was implemented for Doppler imaging. Each acquisition captured one full cardiac cycle (1000 frames; 250 msec). Color and power Doppler images were generated following wall filtering with the SVD (first 50 projections excluded). Doppler spectra were generated from wall filtered data within user-defined regions of interest (ROIs). In brief, the fast Fourier transform was applied to spatially averaged data from the selected ROIs within a sliding temporal window (30 frame window, 2 frame steps). These data were used to derive time varying peak and mean blood flow velocity within a given vessel across a full cardiac cycle. The resistive index was measured as (peak systolic velocity – end diastolic velocity) / peak systolic velocity.

### Immunohistochemistry

After euthanasia, animals were sacrificed via trans-cardiac perfusion with 200 mL of ice-cold PBS (pH 7.4) followed by 200 mL of 4% paraformaldehyde (PFA). Following post-fixation and tissue harvesting, spinal cord tissue was cryoprotected and immersed in optimal cutting temperature medium (Fisher Healthcare Tissue Plus O.C.T. Compound, Clear) before being frozen on dry ice. Spinal segments were sectioned at approximately -20°C using a cryostat (Leica, CM1850). 20 µm thick coronal sections were thaw mounted onto gelatin coated microscope slides and dried before being stored at -80°C.

Fluorescent immunohistochemistry was conducted to analyze microvessel density (laminin or lectin), pericyte coverage (NG2), and microglia density (Iba1). Cryosections were thawed at 37℃ for two hours on a slide warmer followed by a wash with 1x PBS. Sections were then soaked in blocking buffer (3% Normal Goat Serum, 0.3% Triton-X100 in PBS) for one hour before overnight incubation at 4℃ with one of the following primary antibodies: anti-NG2 (1:200; Millipore AB5320), anti-Laminin (Sigma Aldrich, L9393, 1:500), or anti-Iba1 (1:1000, #019-19741, Fujifilm Wako). After washing with PBS, secondary antibody incubation was done using Alexafluor 568 goat anti-rabbit (Invitrogen, A11036, 1:500), Streptavidin Alexa Fluor 488 (Invitrogen, S11223, 1:500), and Alexafluor 488 goat anti-mouse (Invitrogen, A32723, 1:500) at 37℃ for 1hr. Subsequently, sections were incubated in DAPI (1:1000, 10 mins) and mounted with Fluoromount G (Electron Microscopy Sciences, Hatfield, PA).

Individual channel images were captured using a Zeiss AxioZoom microscope and analyzed with ImageJ software. Fluorescence colocalization across all three channels was analyzed to determine the pericyte coverage. The number of pericytes in each vessel and the length of vessels were quantified. Pericyte coverage was calculated as the number of pericytes per length of the microvessel. Microglia density was calculated as the percent area stained positively for Iba1 on binarized images. Researchers were blinded for all measurements.

### Statistical analysis

All summary values are presented as mean ± standard error of the mean (SEM). All statistical analysis and data visualization was conducted using Python (v3.8.10) and the Pandas (v1.4.2), Scipy (v1.8.1), and Seaborn (v.0.11.2) packages. Independent t-tests were used for all comparisons between adult and aged data, and paired t-tests were used to compare normoxic and hypoxic AUC for each condition (α=0.05).

## Results

### Spinal cord volume

Stepped 3D B-mode and 3D CEUS scans were analyzed to identify differences in tissue volume between adult and aged animals. Volumes were measured within a normalized 3 mm rostrocaudal window. B-mode imaging revealed a significantly larger total spinal cord parenchyma volume in aged rats (37.5±0.63 mm^3^ vs 41.9±0.48 mm^3^, p<0.05; Fig. 1A-C). CEUS scans were used to identify more granular differences in gray and white matter (GM, WM) specifically. Higher microvessel density in the GM (3-4x vs WM) in conjunction with CEUS intensity varying proportionally to local microbubble concentration resulted in visible contrast between GM and WM and enabled masking of each tissue component separately (Fig. 1D-E) (9). Aged animals had a significantly higher WM volume, (21.8±0.54 mm^3^ vs 24.9±0.61 mm^3^, p<0.05; Fig. 1F), but not GM (20.6±0.25 mm^3^ vs 21.3±0.24 mm^3^, p>0.05; Fig. 1F), indicating that the increase in total parenchyma volume is attributed solely to WM changes.

**Fig 1.**
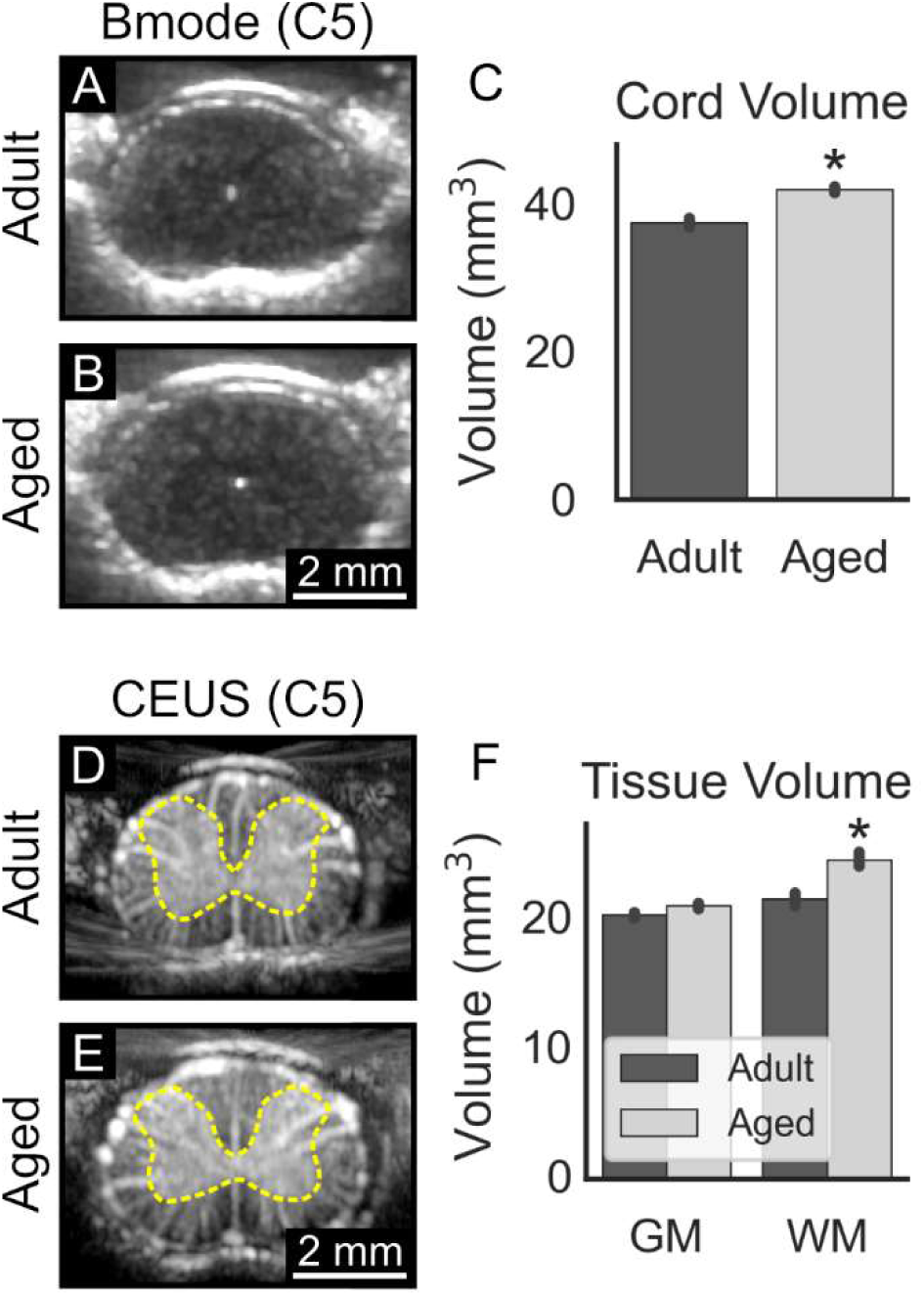
Spinal cord parenchyma volume was significantly larger in aged rats. (A-C) 3D stepped B-mode acquisitions were used to quantify the total spinal cord parenchyma volume within a normalized 3 mm rostrocaudal window centered on C5. Representative axial B-mode images of C5 are shown. Aged animals had significantly larger spinal cords than adult animals (37.5±0.63 mm^3^ vs 41.9±0.48 mm^3^, p<0.05). (D-F) 3D stepped contrast enhanced ultrasound (CEUS) scans were used to visualize and volumetrically measure the white (WM) and gray matter (GM; yellow dashed region). Representative images of C5 are shown. Only the white matter volume was significantly different between adult and aged animals (21.8±0.54 mm^3^ vs 24.9±0.61 mm^3^, p<0.05).

### Vascular resistance

ULM was used to identify differences in vascular anatomy between adult and aged animals. Longitudinal ULM images captured at midline revealed significant tortuosity in the anterior spinal artery (ASA; Fig. 2A-B, white arrows) of aged animals (tortuosity index 2.20±0.15 vs 4.74±0.45, p<0.05; Fig. 2C). Significant tortuosity in the major feeder artery for the observed segments (C5-C6, Fig. 2) likely contributed to elevated vascular resistance observed with both CEUS bolus and Doppler imaging. CEUS bolus imaging revealed significantly higher rise time specifically in the microvasculature in the aged cohort (1.73±0.17 sec vs 2.26±0.07 sec, p<0.05; Fig. 2D-F). Spectral Doppler imaging identified a significantly higher resistive index in aged rats, specifically in the central sulcal arteries (CSAs) that branch from the ASA and supply blood to the spinal cord (0.37±0.02 vs 0.43±0.02, p<0.05). Collectively, these data indicate significantly elevated vascular resistance in the aged cohort.

**Fig 2.**
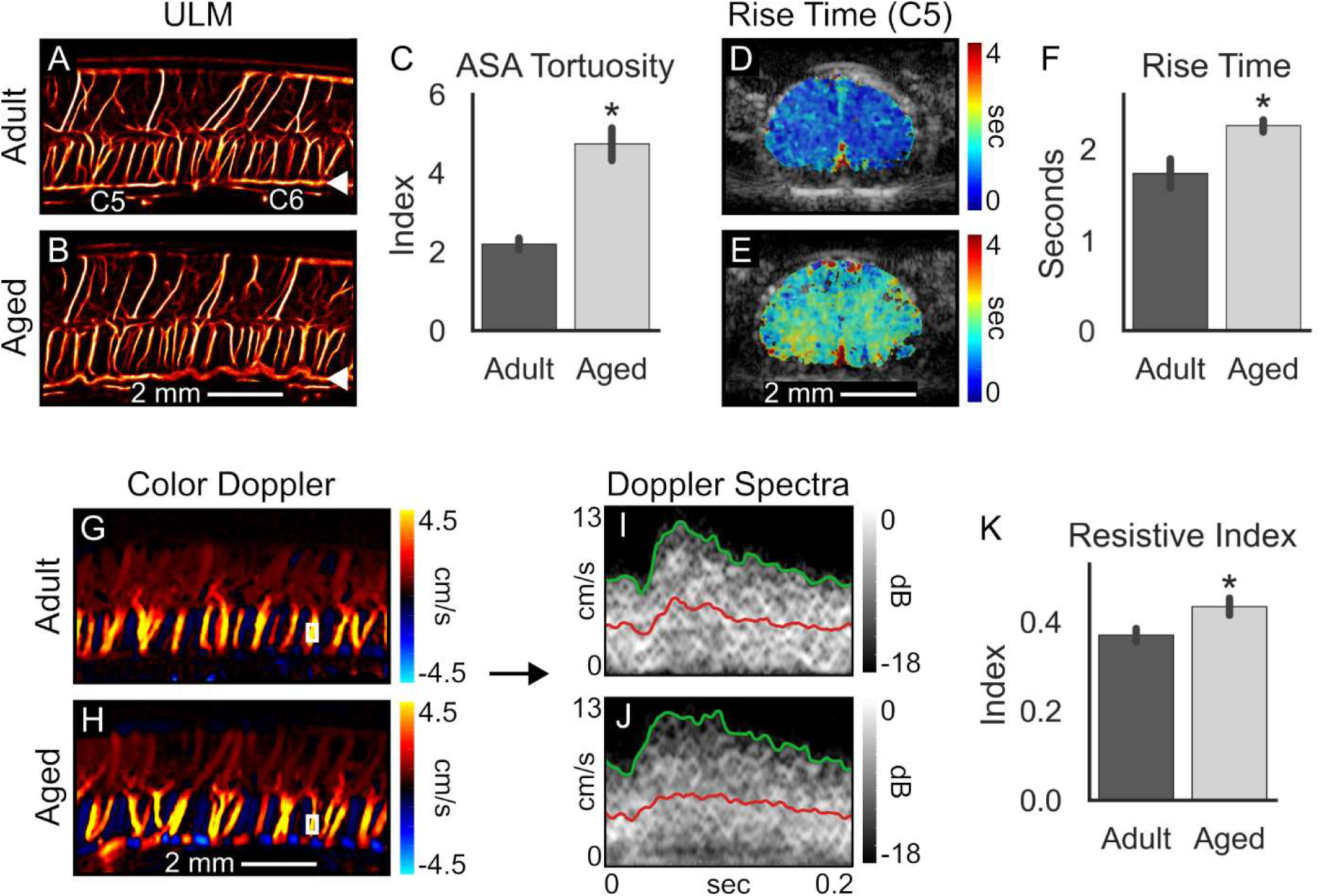
Baseline vascular resistance is elevated in aged rats. (A-C) Ultrasound localization microscopy (ULM) was used to visualize the spinal vasculature at high resolution *in vivo*. Representative longitudinal images acquired at C5-C6 are shown. Significantly elevated tortuosity was observed in the anterior spinal artery (ASA; white arrow) in aged animals (tortuosity index 2.20±0.15 vs 4.74±0.45, p<0.05). (D-F) Rise time, a metric analogous to vascular resistance in the microcirculation, was derived from bolus inflow acquisitions. Example cross-sectional images acquired at C5 are displayed. Aged animals exhibited significantly elevated rise time (1.73±0.17 sec vs 2.26±0.07 sec, p<0.05). (G-H) Longitudinal color Doppler images are displayed for representative adult and aged animals. (I-K) Spectral Doppler analysis was used to quantify the resistive index, a measure of resistance to flow, in the central sulcal arteries (CSAs). Representative spectra for a selected vessel (white rectangle in G-H) are shown for a single cardiac cycle. Mean flow velocity is traced in red; peak velocity is traced in green. The resistive index was significantly elevated in aged rats (0.37±0.02 vs 0.43±0.02, p<0.05).

### Blood volume and microvessel density

Power Doppler and CEUS bolus imaging were used to identify baseline differences in blood volume between adult and aged animals. The power Doppler signal intensity has been shown to vary proportionately to local blood volume; this is the underlying foundation of functional ultrasound imaging and has been extensively reported (23, 24). Here, CSAs and dorsal ascending veins were independently analyzed using manually placed ROIs. Aged animals exhibited a significantly lower Doppler signal intensity, and by extension a lower blood volume, in both CSAs and veins (CSA: 2.08e14±1.62e13 vs 1.46e14±1.05e13, p<0.05; vein: 4.91e14±4.65e13 vs 3.13e14±3.18e13, p<0.05; Fig. 3A-C). CEUS bolus data revealed significantly reduced blood volume in the microcirculation specifically in WM (4.44e14±1.37e13 vs 3.66e14±2.64e13, p<0.05) but not in GM (7.77e14±3.24e13 vs 7.23e14±3.41e13, p>0.05; Fig. 3D-F). Immunohistochemistry corroborated these findings; analysis of tissue sections stained for laminin (pan-vascular marker) showed a significantly lower microvessel density (MVD) in WM (84.2±2.46 vs 67.4±0.63 vessels/mm^2^, p<0.05) but not GM (427±26.5 vs 424±15.5 vessels/mm^2^, p>0.05; Fig. 3G-I).

**Fig 3.**
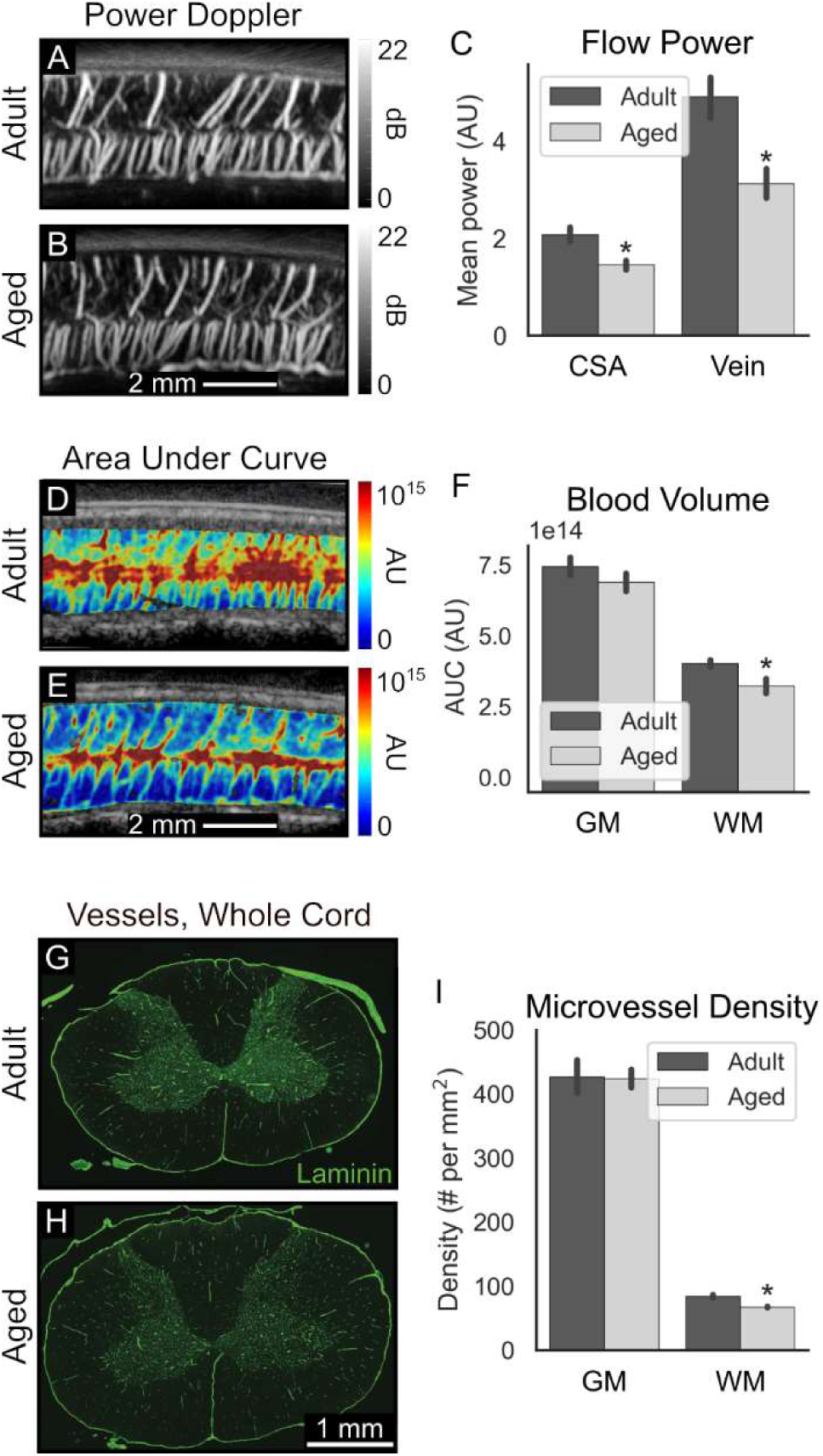
Baseline blood volume is reduced in aged animals only in white matter. (A-C) Power Doppler imaging was conducted to measure flow power (analogous to blood volume) in the dorsal ascending veins and central sulcal arteries (CSAs). Aged animals exhibited significantly lower flow power than adult animals (3.49e14±6.90e13 vs 2.29e14±4.21e13, p<0.05). (D-F) The area under the curve (AUC), derived from bolus inflow acquisitions, was used as a relative measure of local blood volume in the microcirculation. Aged animals exhibited significantly lower baseline blood volume specifically in the white matter (4.44e14±1.37e13 vs 3.66e14±2.64e13, p<0.05). *In vivo* findings were corroborated with immunohistochemistry. (G-H) Whole cord, cross-sectional images of laminin-stained sections (vessels; green) are displayed. (I) Aged rats had significantly lower microvessel density in the white matter (84.2±2.46 vs 67.4±0.63 vessels/mm^2^, p<0.05) but not in the gray matter (p>0.05).

### Hypoxia challenge and potential cellular mediators

CEUS bolus imaging was conducted under normoxic (20% O_2_) and hypoxic (10% O_2_) conditions to identify how adult and aged animals respond to a vascular challenge. Adult animals exhibited a significant hyperemic response during hypoxia in both GM and WM (GM: 1.13±0.10-fold change from normoxia, p<0.05; WM: 1.16±0.13, p<0.05). Aged animals showed no hemodynamic response during hypoxia (GM: 1.00±0.08; WM: 1.03±0.08, p>0.05; Fig. 4A-E). Immunohistochemical analysis was used to identify potential cellular mediating factors for the lack of response in the aged cohort. Specifically, the prevalence of pericytes and microglia was quantified; both cell types have previously been implicated in regulating neurovascular response. Analysis revealed significantly lower pericyte coverage in WM in aged animals (0.017±0.002 vs 0.007±0.002 cells per µm vessel length, p<0.05; Fig. 4F-J) and significantly higher microglia density in the GM in aged animals (0.65±0.11% vs 1.24±0.13% area coverage, p<0.05; Fig. 5).

**Fig 4.**
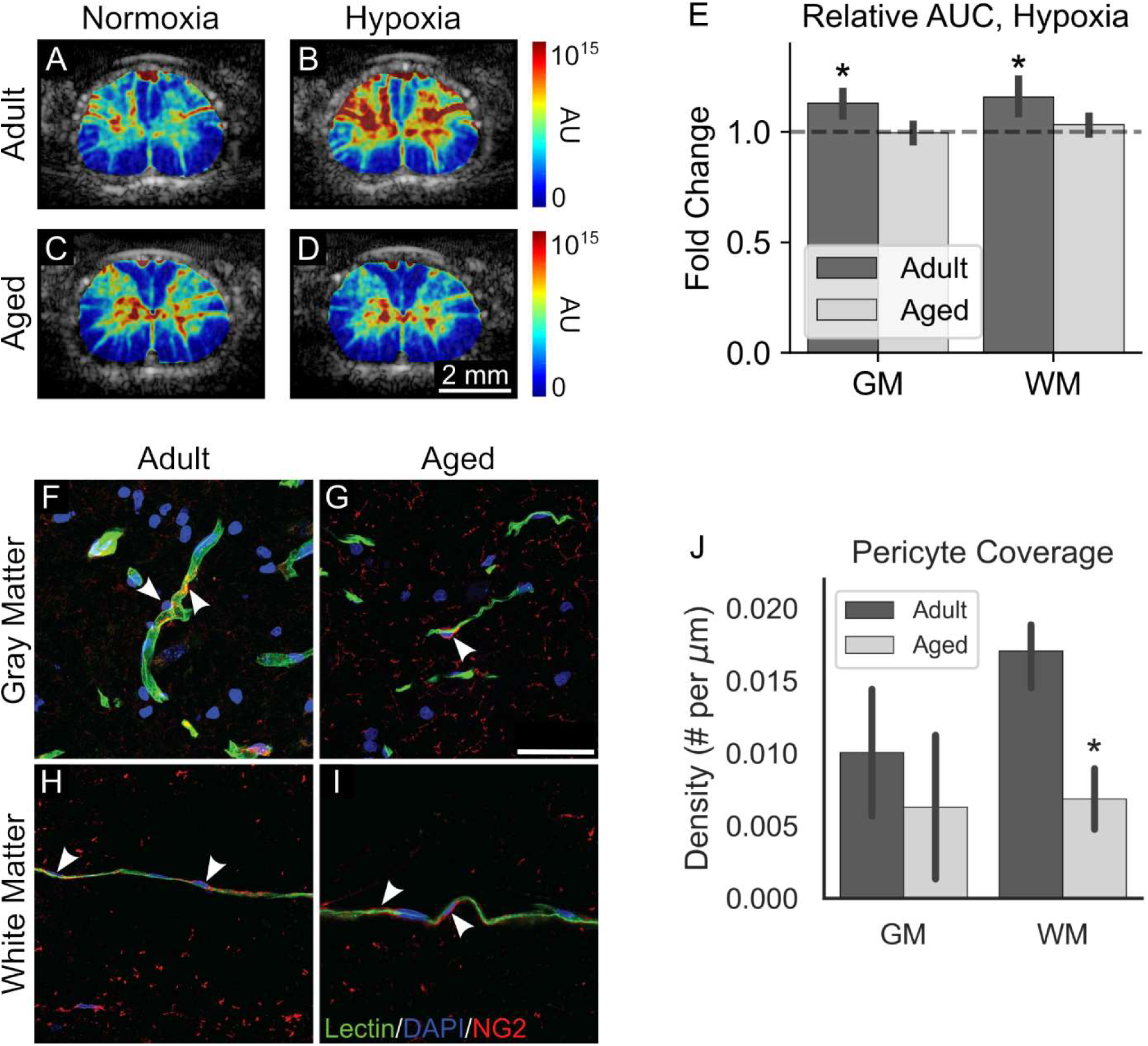
Aged rats did not respond to hypoxia. Area under the curve (AUC), derived from bolus inflow acquisitions, was used to quantify relative blood volume before (normoxia) and during a hypoxia challenge (10% O2, 3 min on). (A-D) Representative cross sectional AUC maps acquired at C5 are shown. (E) While adult rats showed significantly elevated blood volume in both gray and white matter during hypoxia (GM: 1.13±0.10-fold change from normoxia, p<0.05; WM: 1.16±0.13, p<0.05), aged rats showed no response (GM: 1.00±0.08; WM: 1.03±0.08). Pericytes were investigated to determine their role in the differential response to hypoxia between adult and aged rats. (F-I) Representative images depicting sections stained for lectin (vessels; green), DAPI (nuclei; blue), and NG2 (pericytes; red) are shown. Example vascular-associated pericytes and their projections are highlighted with white arrows. (J) Pericyte coverage, measured as the number of pericytes per unit vessel length, was significantly reduced in the white matter in aged animals (0.017±0.002 vs 0.007±0.002, p<0.05).

**Fig 5.**
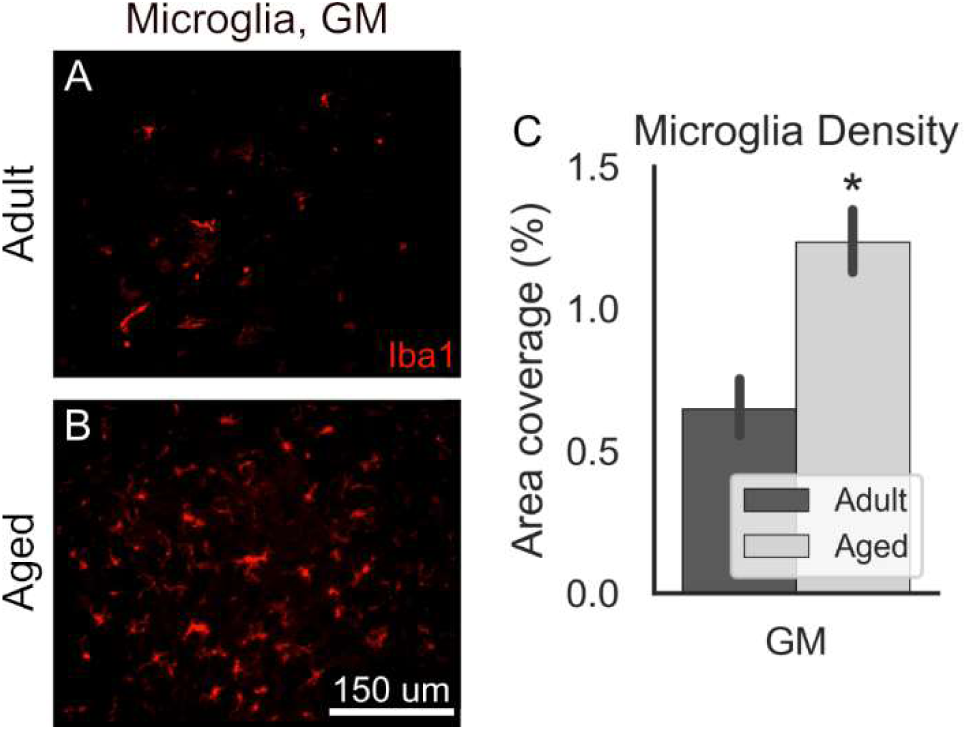
Microglia density is higher in the gray matter in aged rats. (A-B) Representative images of Iba1-stained sections (microglia; red), zoomed to focus on the gray matter, are shown. (C) Microglia density was significantly higher in the gray matter in aged rats (0.65±0.11% vs 1.24±0.13%, p<0.05).

## Discussion

This study provides the first *in vivo* evidence of both hemodynamic and anatomical vascular alterations in the rat spinal cord with normal aging. To date, most *in vivo* studies of spinal hemodynamics have relied on depth-limited optical imaging modalities (e.g., laser speckle contrast, two photon imaging) (12, 13). These are incapable of visualizing the full dorsal-ventral extent of the spinal cord and have had limited utility in examining differences in gray and white matter. Ultrafast CEUS imaging is not subject to the same depth limitations and has extended the capabilities of these existing modalities by enabling segmentation and independent analysis of flow at different levels of the vasculature (16, 19). We applied these segmentation capabilities to look at baseline differences in vascular resistance (Fig. 2) and blood volume (Fig. 3) in both larger vessels and microvasculature, and to investigate microvascular-specific responses to a hypoxia challenge (Fig. 4). Moreover, to our knowledge, there have been no prior *in vivo* studies investigating hemodynamic differences in the spinal cord due specifically to normal aging. A single study was conducted using standard *ex vivo* methods to quantify differences in vascular anatomy (9). Another study examined vascular disruption and impaired angiogenesis as potential mediating factors for worse functional outcomes in aged rodents following spinal cord injury, though this study also relied solely on *ex vivo* methods (11). The current study provides a basis for further work elucidating the role of spinal microvascular alterations in normal aging and potentially in enhancing susceptibility to ischemic injury with age.

Volumetric CEUS analysis revealed significantly elevated WM volume in aged rats (Fig. 1). There is some evidence demonstrating that the aged spinal cord exhibits axonal loss and demyelination, similar to the aging brain (25–27). This process is likely associated with significantly elevated microglia density, particularly microglia with morphological changes characteristic of a reactive phenotype, in the aged WM (28). One study found a number of cystic regions within the WM associated with elevated staining for glial fibrillary acidic protein (GFAP) and an increased prevalence of Marchi positive bodies in aged rats (29). This progressive degenerative process and in particular the formation of cystic regions may contribute to the observed increase specifically in WM volume in the current study. We also found both a slight decrease in MVD and a significant decrease in baseline blood volume specifically in the white matter (Fig. 3), indicating that the WM microvasculature is also impaired due to normal aging. Further work is necessary to fully reconcile the changes in WM morphology and vascular function with the known progressive functional degeneration in WM in aged rats. One target for future investigation is blood spinal cord barrier breakdown with normal aging and associated edema; while blood brain barrier degradation has been reported with age, less is currently known in the spinal cord (30, 31). Existing optical techniques struggle in the spinal cord due to increased scattering through heavily myelinated white matter; CEUS is not subject to these limitations, and therefore represents a uniquely useful tool to quantify white matter differences in these future studies (14, 15).

Elevated vascular resistance in both penetrating central sulcal arteries and the microcirculation was observed in conjunction with significant tortuosity in the anterior spinal artery in the aged cohort (Fig. 2). One prior study showed elevated tortuosity in the spinal vasculature of aged rats, as well as vessel dropout and reduced microvessel density, but no prior studies have demonstrated hemodynamic differences in the spinal cord (9). Several studies have elucidated age-related hemodynamic changes in the brain, however. Progressive increases in vascular resistance have previously been reported in aged patients, particularly those exhibiting cognitive decline (32, 33). Increased baseline vascular resistance reduces the ability to respond to metabolic demand and can impair neurovascular coupling (34). Dysregulation of control over vascular tone may also contribute to either hypoperfusion in the case of rapid blood pressure drop, or elevated blood pressure and resulting damage in the capillary network due to reduced buffering capacity in the upstream arteries (34). Collectively, these changes can contribute to cognitive decline and the progression of neurodegenerative diseases (32, 33, 35). Based on the findings from the current study, the aged spinal cord may be similarly impacted.

A hypoxia challenge was administered to elucidate functional microvascular changes in the aged spinal cord. While adult animals exhibited a significant hyperemic response during hypoxia, aged animals did not respond at all (Fig. 4). While therapeutic intermittent hypoxia exposure and associated induced neuroplasticity in the spinal cord is well-described, little work has been done to elucidate associated spinal hemodynamic changes even in healthy adults (36, 37). A substantial amount of previous work has investigated changes in cerebral blood flow in response to hypoxia challenge in patients and in preclinical models. Healthy adults consistently exhibit a hyperemic response to account for increased oxygen demand under hypoxic conditions (38, 39). Normal aging has previously been implicated in impaired cerebrovascular response to hypoxia, corroborating the findings from the current study (40, 41).

Recently, both pericytes and glial cells, including microglia, have been implicated as mediating cellular factors in the regulation of cerebral blood flow (42–44). The current study found a reduction in pericyte coverage and dysregulated microglia expression in the aged spinal cord, suggesting two possible contributing factors to the lack of microvascular response under hypoxia exposure. The inability to respond to increased metabolic demand may contribute to mild hypoxia under stress conditions with normal aging, increased susceptibility to ischemic injuries and impaired functional recovery, and may be implicated in the early stages of certain neurodegenerative disorders (32, 34, 45–47). Our lab has recently used CEUS imaging to quantify secondary spinal cord injury expansion in healthy adult rodents; future work will apply these techniques to investigate differential injury expansion in an aging model (17).

One limitation of the current study is the use of a single measurement during hypoxia. More complex temporal changes may be occurring during hypoxia exposure, particularly in the aged rats. Future work will utilize functional ultrasound (fUS) recordings, which afford substantially more granular temporal information and provide additional quantitative features for comparison (e.g., onset time, peak response, decay rate, etc.) (24). The current study is also limited by the subject demographics; we only used male rats, and we only examined adult and aged animals. Sex is a critically relevant biological variable for the response to spinal cord injury in normal adults and has been implicated in differences in spinal cord inflammation and gene expression with normal aging (48, 49). Age-dependent changes may occur in a progressive or stepwise manner, necessitating further study with young, middle aged, and aged cohorts. The imaging techniques described here may also be applicable for a number of relevant age-associated pathologies, including spinal cord injury (as previously described), and progressive cervical myelopathy (50).

## Conclusion

CEUS imaging revealed significant anatomical and functional alterations in the aged spinal cord. In addition to reduced blood volume and elevated vascular resistance at baseline, aged rats failed to respond to a hypoxia challenge. Future work will broaden the study population to include animals from both sexes and more granular age groups and will investigate the impact of different pathologies (spinal cord injury, cervical myelopathy) on blood flow in the aged spinal cord.

## Acknowledgments

Author contributions: **JNH**: Methodology, Software, Investigation, Writing – Original Draft, Writing – Review and Editing, Visualization. **PC**: Investigation, Writing – Review and Editing, Visualization. **AC**: Writing – Review and Editing, Investigation, Visualization. **JEH**: Writing – Review and Editing, Investigation, Visualization. **GJH**: Writing – Review and Editing, Investigation. **MJR**: Conceptualization, Funding Acquisition, Writing – Review and Editing. **MFB**: Supervision, Writing – Review and Editing. **ZZK**: Conceptualization, Methodology, Investigation, Writing – Review and Editing, Supervision, Project Administration, Funding Acquisition.

